# Somatic cells non-autonomously control germline incomplete cytokinesis through FGF signaling

**DOI:** 10.64898/2026.06.04.729929

**Authors:** Beth Kern, Zachary Y. Berkley, Samuel Price, Kari F. Lenhart

**Affiliations:** Biology Department, Drexel University

**Keywords:** FGF, F-actin, cytokinesis, differentiation

## Abstract

Across species, germ cells divide and differentiate as interconnected units, termed cysts. These cysts are generated through reiterative rounds of mitosis followed by incomplete cytokinesis to generate stable ring canals (RCs). Despite the ubiquity of germ cell incomplete cytokinesis, it is still unclear how this program is mechanistically regulated across multiple cell cycles to retain integrity of the cyst. Here, by leveraging longitudinal live imaging of the *Drosophila* testis we have identified a critical, non-autonomous role for somatic support cells in maintenance of germline RC stability. We find that F-actin at RCs is stable throughout interphase but is dynamically disassembled and reassembled at each reiterative mitotic entrance and exit. Importantly, we find that somatic cells regulate the stability of interphase RC F-actin through the secreted growth factor, FGF. Genetic or pharmacological inhibition of FGF signaling induces disassembly of RC F-actin during interphase. Persistent clearance of F-actin from the RC leads to failure of incomplete cytokinesis and cyst abscission, suggesting that stable F-actin at RCs is required for the robust maintenance of incomplete cytokinesis through multiple rounds of germ cell divisions. Finally, we mechanistically link FGF signaling to germline activity of the non-receptor tyrosine kinase, Src64, which is known to regulate RC F-actin through Arp2/3. Taken together, we find a previously unappreciated role for somatic support cells in controlling an essential aspect of germ cell biology in the mitotically dividing spermatogonial pool.

**Summary Statement:** Somatic cells of the gonad secrete FGF ligand, Pyramus, which is required for maintenance of F-actin at germline ring canals and integrity of germline incomplete cytokinesis.

## Introduction

Adult stem cells promote tissue homeostasis through consistent production of specialized progenitor cells. Yet constant cell divisions are deleterious, increasing the chances for genomic instability and subsequent tissue degeneration (Yang & Yamashita, 2015; Zhang & Hsu, 2017). To preserve the stem cell pool, the major mitotic burden is placed on progenitor cells, which undergo extensive mitotic expansion during differentiation (Zhang & Hsu, 2017). This “transit-amplification” (TA) of progenitor cells is required for homeostasis across multiple tissues, ranging from the eye to the intestine (Barker, 2014; Lavker & Sun, 2003).

In no tissues are TA divisions more critical than the male and female germline. Across species, differentiating germ cells engage a specialized and highly conserved TA program in which mitotic divisions occur with incomplete cytokinesis. Ultimately, this generates germ cell cysts that retain connectivity through stable intercellular bridges (ICBs), termed ring canals (RCs) (Fig.1A) (Greenbaum et al., 2006; Price et al., 2023). In both males and females, formation of stable RCs is essential for proper germ cell differentiation. During oogenesis, RCs allow for cytoplasmic sharing and eventual transfer of RNAs, proteins, and organelles into the oocyte (Lei & Spradling, 2016; K. L. Lu & Yamashita, 2017). In males, germ cell interconnectivity promotes robust detection and elimination of genomically damaged germ cells prior to meiosis (K. L. Lu & Yamashita, 2017). Thus, generation and maintenance of RCs through incomplete cytokinesis is essential for reproduction in both sexes and across species.

**Fig. 1.**
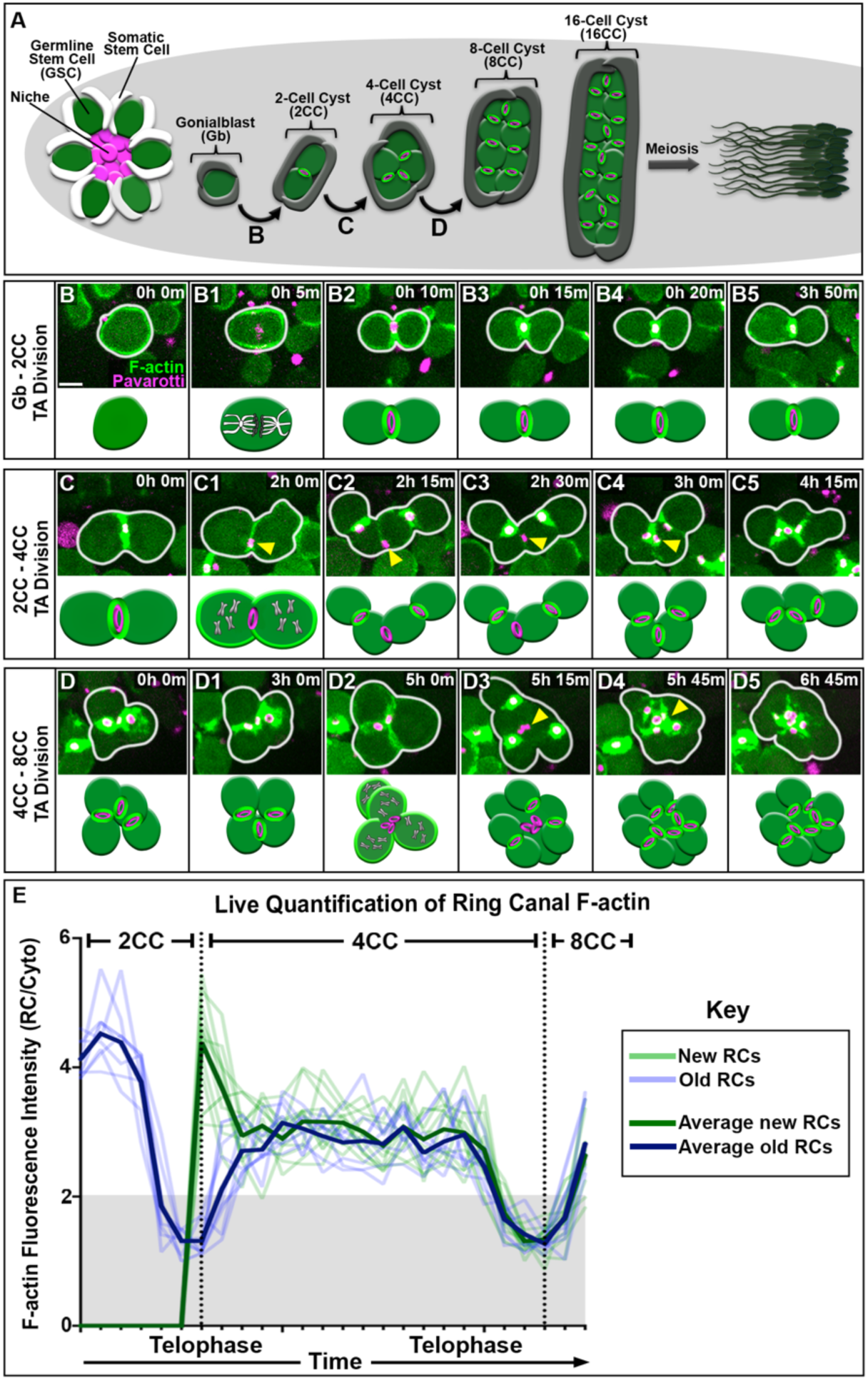
Dynamic regulation of RC F-actin during TA divisions. (A) Diagram of the *Drosophila* testis. (B-D) Time-lapse imaging of nos-ABD-moe::GFP, Pav::mCh and associated diagrams (combined 1-3D TjGal4^TS^ and nosGal4 controls) through three rounds of germ cell cyst mitosis and incomplete cytokinesis. (B) Gb-2CC division (n=9). (C) 2CC-4CC division (n=83). (D) 4CC-8CC division (n=87). Each image is 1-6 z-slices to capture the division plane. Images acquired every 15 minutes. (E) Quantification of normalized RC F-actin intensity over multiple rounds of germ cell cyst divisions (n=8).

In most organisms, RCs consist of an internal ring derived from the cytokinetic midbody surrounded by a dense F-actin network (Gary R. Hime et al., 1996; Haglund et al., 2011; Price et al., 2023). Given these structural and molecular components, RCs have been described as “arrested” contractile rings (CRs) with the assumption that RC genesis occurs via arrest of the traditional cytokinetic program (Gary R. Hime et al., 1996; Haglund et al., 2011; Price et al., 2023). Recently, live imaging has challenged this belief by showing that the compacted cytokinetic midbody undergoes a regulated expansion event to generate the RC and retain interconnectivity with cysts (Price et al., 2023). These findings support the notion that RC genesis is not simply arrested cytokinetic progression but is instead a dynamic and highly regulated event.

Indeed, many of the proteins identified to play a role in RC stability do so independently of the canonical cytokinesis program. In mice, the kinase testis-expressed gene 14 (TEX14) localizes to RCs and directly inhibits abscission by preventing recruitment of ALIX and ESCRT I component TSG101 (Greenbaum et al., 2006, 2011; Iwamori et al., 2010). In *Drosophila*, Src64 kinase controls RC integrity via F-actin polymerization by regulating the crosslinker Kelch in females and activating an Abelson Kinase (Abl)-Arp2/3 cascade in males (Cooley, 1998; Eikenes et al., 2015; Robinson & Cooley, 1996). In both sexes, disruption of Src64 causes defects in RC morphology and germ cell differentiation. Interestingly, the kinase domain in mammalian TEX14 resembles that of Src64 (Greenbaum et al., 2006), suggesting possible conservation across species in regulation of RC stability. However, the upstream signals activating these kinases to control incomplete cytokinesis are entirely unknown.

In all species, the germline is intimately associated with somatic support cells (Fairchild et al., 2015; John Tran et al., 2000; Le Bras & Van Doren, 2006; Lim & Fuller, 2012; Masaki et al., 2018; Oatley & Brinster, 2012; Takashima et al., 2015). Somatic cells adhere to and regulate several essential aspects of germline biology, including germline stem cell cytokinesis (Lenhart & DiNardo, 2015), progenitor differentiation (Gonczy et al., 1997; Insco et al., 2009; Kawase et al., 2004; C. Y. Li et al., 2007; Tang et al., 2017a), and transit amplifying division “count” (Insco et al., 2009). Yet a role for somatic support cells in mediating the highly conserved incomplete cytokinesis within germline cysts has never been explored.

Here, we used a combination of genetics and live imaging to identify a non-autonomous role for somatic in successful execution of incomplete cytokinesis within the germline. Critically, we find that maintenance of F-actin at RCs requires activation of the conserved RTK, FGFR, in germ cells by a soma-secreted FGF ligand. Depletion of FGF ligand Pyramus (Pyr) in somatic cells, FGFR Heartless (Htl) in the germline or pharmacological inhibition of FGFR activity disrupts F-actin stability at RCs and permits inappropriate completion of cytokinesis in germline cysts. Immunofluorescent and genetic data establish a direct mechanistic link between FGF signaling and activation of the known Src64-Abl-Arp2/3 pathway previously shown to regulate RC F-actin (Eikenes et al., 2015). Taken together, our data identify a novel role for somatic cells in regulating the highly conserved incomplete cytokinesis program within the germline.

## Results

### Dynamic regulation of RC F-actin over the germline cyst cell cycle

To investigate RC dynamics across multiple rounds of germ cell TA divisions, we performed 24-hour time-lapse imaging of adult testes, observing up to three rounds of mitosis within a single cyst (Fig.1A) (Lenhart & DiNardo, 2015; Roach & Lenhart, 2024). To examine incomplete cytokinesis, we visualized F-actin in germ cells via expression of the actin binding domain of moesin fused to GFP (nos-ABD-moe::GFP), and midbody rings by an endogenously tagged Pavarotti::mCherry (Pav::mCh) (Price et al., 2023).

Germline stem cells (GSCs) divide and generate a single daughter cell, the gonialblast (Gb). During the first TA division (Fig.1B, MovieS1), a Gb enters mitosis (Fig.1B1), assembles an F-actin CR (Fig.1B3), and persistently maintains F-actin from the CR at the nascent RC following mitotic exit (Fig.1B3-Fig.1C). This produces a 2-cell cyst (2CC) with a stable RC. However, F-actin at RCs is not permanently stabilized.

Indeed, there is rapid F-actin disassembly at the existing RC as the 2CC enters the next round of TA division (Fig.1C1, arrowhead; MovieS2). As mitosis progresses, acto-myosin CRs are generated and stabilized between nascent daughter cells of the 4CC, while F-actin remains absent from the existing “old” RC (Fig.1C2-C3, arrowhead). F-actin is absent from the old RC for an average of 45 minutes during this 2CC-4CC division. It is only after formation of new RCs and mitotic exit of the 4CC that the old RC undergoes a distinct, *de novo* F-actin reassembly event (Fig.1C4, arrowhead). All three RCs, once assembled, maintain F-actin for the duration of the 4CC interphase (Fig.1C4-D1). However, upon entry into another round of mitosis, F-actin is depolymerized at all existing RCs (Fig.1D2; MovieS3). Four new RCs are generated, all of which maintain CR F-actin (Fig.1D3; MovieS3). Again, F-actin remains absent from the three oldest RCs until eventual re-assembly (Fig.1D3-D4, arrowheads; MovieS3).

To quantify the dynamics of this process, we identified germ cell cysts that executed two rounds of TA division during a single imaging period and quantified the intensity of RC F-actin from the 2CC-8CC stages (Fig.1E). By temporally aligning quantification of each cyst to mitotic telophase, we examined population-level trends in F-actin dynamics over time and across multiple cell divisions. To establish a quantitative metric for F-actin presence or absence at the RC, we analyzed F-actin during complete cytokinesis in GSCs. By comparing the mean intensities of F-actin at the ICB during early cytokinesis (F-actin present) versus late cytokinesis (F-actin absent), we calculated a 2-fold or higher enrichment of F-actin relative to the germ cell cytoplasm denotes “presence” of F-actin at the ICB (Fig.S1A-1B). Thus, any RC F-actin ratio below 2 indicates an absence of F-actin at that RC (Fig.1E, grey rectangle). Consistent with our qualitative analyses, live quantification of RC F-actin intensities revealed dramatic disassembly of F-actin immediately prior to mitotic entry followed by rapid reassembly of F-actin following mitotic exit. All RCs, both new and old, then retain F-actin for the duration of interphase (Fig.1E). This dynamic disassembly and reassembly of F-actin at old RCs is repeated at the next TA division (Fig.1E). Taken together, our live imaging reveals a dynamic and tightly regulated process by which F-actin at RCs is reiteratively reestablished during TA divisions of germline cysts.

### FGF ligand Pyramus is expressed by somatic cells and activates FGFR in the germline

Given the importance of somatic support cells in regulating germline biology (Ikami et al., 2015; Kawase et al., 2004; Masaki et al., 2018; Takashima et al., 2015; Tang et al., 2017b), we asked whether soma-germline signaling might also control F-actin dynamics and RC integrity. To identify candidate somatic signals, we mined transcriptomic data from single cell RNA sequencing (Hof-Michel & Bökel, 2020; Raz et al., 2023). In both datasets, expression of the FGF ligand Pyramus (Pyr) is enriched in differentiating somatic cells, while germ cells express the FGF receptor Heartless (Htl) (Fig.S1A,B). Given the known roles for FGF in regulating F-actin dynamics (Kadam et al., 2009; J. Li et al., 2025; Olson & Nordheim, 2010) and mammalian germ cells (Bowles et al., 2010; Masaki et al., 2018; Takashima et al., 2015), we interrogated a potential role for somatic FGF signaling in germline incomplete cytokinesis.

In accordance with RNA sequencing data, we find that Pyr is expressed in the testis and predominantly colocalizes with somatic cell membranes (Fig.2A-B’’) (Stepanik et al., 2020). Somatic expression of *pyr* RNAi induced a significant reduction in Pyr fluorescence intensity at the testis apex, confirming antibody specificity (Fig.2C-E). In addition, a well-characterized Gal4 line under control of *pyr* regulatory sequences drives expression of nuclear reporter Red Stringer in somatic cells (Fig.S2A). Thus, the FGF ligand Pyr is indeed expressed and secreted by somatic cells of the testis.

**Fig 2.**
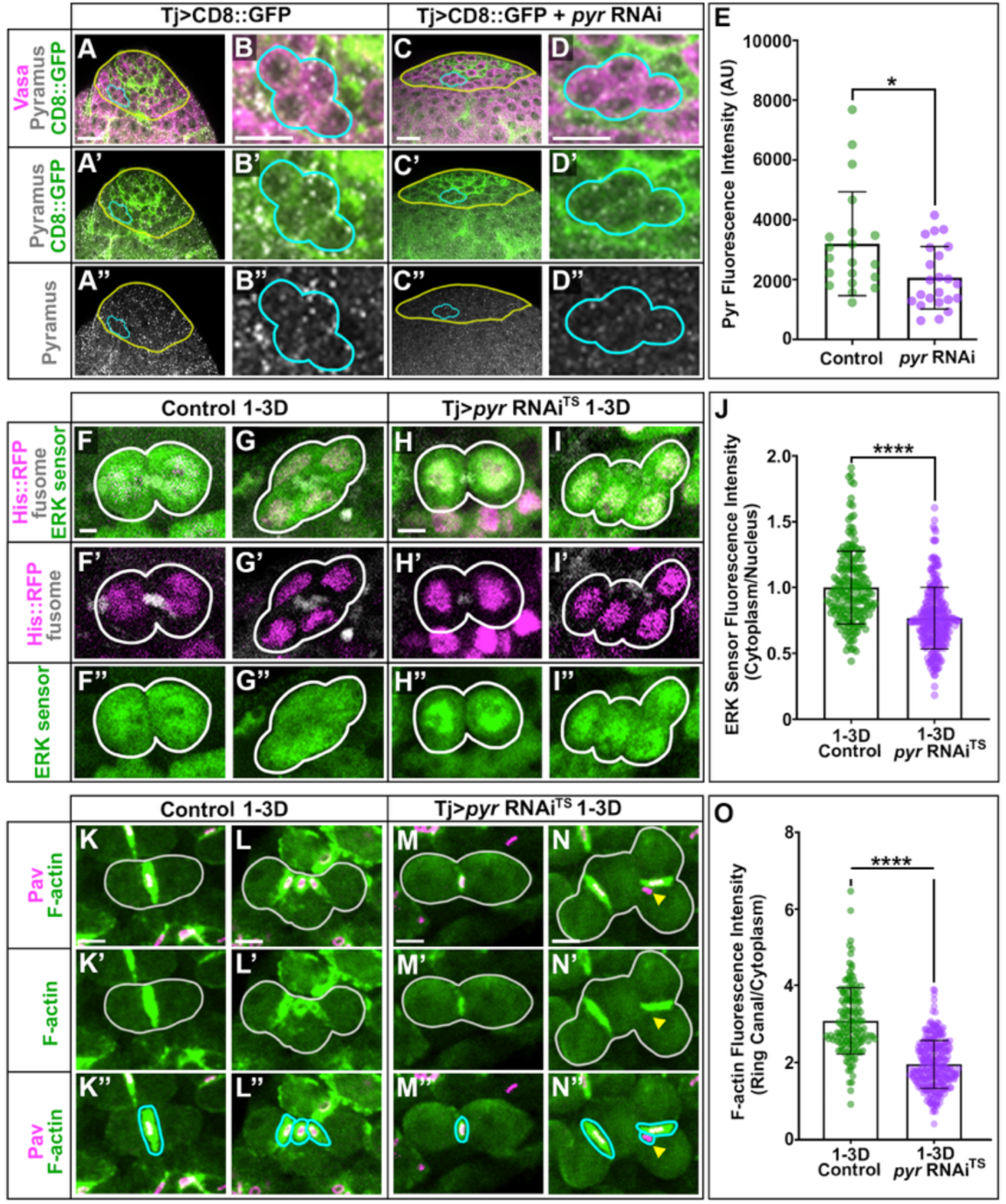
FGF ligand Pyramus is expressed by somatic cells and activates FGFR in the germline. (A-A”, C-C”) Testis apical tip in Tj>CD8::GFP (A-A”) and Tj>CD8::GFP, *pyr* RNAi (C-C”). Regions of interest outlined in yellow. Scale = 40µm. (B-B”, D-D”) Representative 4CC in control (B-B”) and *pyr* RNAi (D-D”) testes. Scale = 5µm. (E) Pyr mean fluorescence intensity (control n=20; pyr RNAi, n= 23) (Mann Whitney t-test p=0.0154). (F-I”) Representative 2CCs and 4CCs (Erk-KTR::Clover, His::mCh, fusome in gray) in (F-F”, G-G”) 1-3D Tj^TS^ control and (H-H”, I-I”) Tj>*pyr* RNAi^TS^. Scale = 5 µm. (J) Quantification of cytoplasmic to nuclear fluorescence intensity of the Erk sensor in control (n=186) and *pyr* RNAi^TS^ (n=256) (p<0.0001, Mann Whitney t-test). (K-N”) F-actin in representative 2CCs and 4CCs (nos-ABD-moe::GFP, Pav::mCh) in (K-K”, L-L”) control 1-3D TjGal4^TS^ and (M-M”, N-N”) 1-3D Tj>*pyr* RNAi^TS^. (O) Quantification of RC F-actin intensity in fixed testes (control n=150, *pyr* RNAi^TS^ n=274. p<0.0001, Mann Whitney t-test). Scale = 5 µm.

To confirm that somatic Pyr signals to the adjacent germline, we utilized an established activity sensor for the FGFR intracellular effector Erk (Wilcockson et al., 2023; Yuen et al., 2022). This GFP reporter consists of a docking site for Erk fused to nuclear import and export sequences, so that pathway activity can be assessed by measuring the ratio of cytoplasmic to nuclear fluorescence. Using expression of a temperature-sensitive Gal80, we expressed *pyr* RNAi in somatic cells (Tj>*pyr* RNAi^TS^ or *pyr* RNAi^TS^) for 1-3 days and assessed FGF pathway activity in the germline. Strikingly, somatic depletion of *pyr* was sufficient to significantly reduce germ cell Erk activity (Fig.2F-J). We observe a similar result with expression of RNAi against FGFR Htl in germ cells (Fig.S2D-H). Taken together, we find that just as in mammals, somatic cells of the *Drosophila* testis secrete FGF, activating FGFR in adjacent germ cells.

### Somatic cells control F-actin stability at germline RCs via FGF signaling

To interrogate if FGFR activation regulates incomplete cytokinesis, we first quantified F-actin intensity at RCs in fixed testes of control and Tj>*pyr* RNAi^TS^ flies (Fig.2K-O). Strikingly, we found that inhibition of FGF signaling significantly reduced RC F-actin in both 2CCs and 4CCs (Fig.2O). In controls, F-actin intensity at RCs was ∼3-fold higher than the germ cell cytoplasm (Fig.2K-2L”, 2O). By contrast, somatic depletion of *pyr* reduced F-actin enrichment at RCs to only 1.97 (Fig.2M-2O). Critically, this average is just below the calculated threshold for F-actin presence at ICBs, indicating FGF inhibition results in significant depolymerization of F-actin at RCs.

We next performed live imaging of F-actin dynamics upon FGF pathway inhibition. As expected, control RCs maintain F-actin for the majority of the 4CC cell cycle, only disassembling F-actin during mitosis and then rapidly reestablishing F-actin during interphase (Fig.3A-A5). By contrast, both Tj>*pyr*RNAi^TS^ and *nos*>*htl* RNAi testes exhibited defects in RC integrity over time. In both genetic backgrounds, transit amplification was successful until the 4CC stage. However, loss of FGF signaling consistently led to failure of F-actin maintenance at RCs across the 4CC interphase (arrowheads, Fig.3B1-B5 and C1-C5). These F-actin defects ranged from mild, in which RC F-actin dynamically disassembles several times across the cell cycle but was ultimately restored, to severe, in which RC F-actin depolymerized and remained persistently absent. Critically, persistent F-actin absence frequently led to inappropriate completion of cytokinesis (Fig.3B5, C5, MovieS4).

**Fig. 3.**
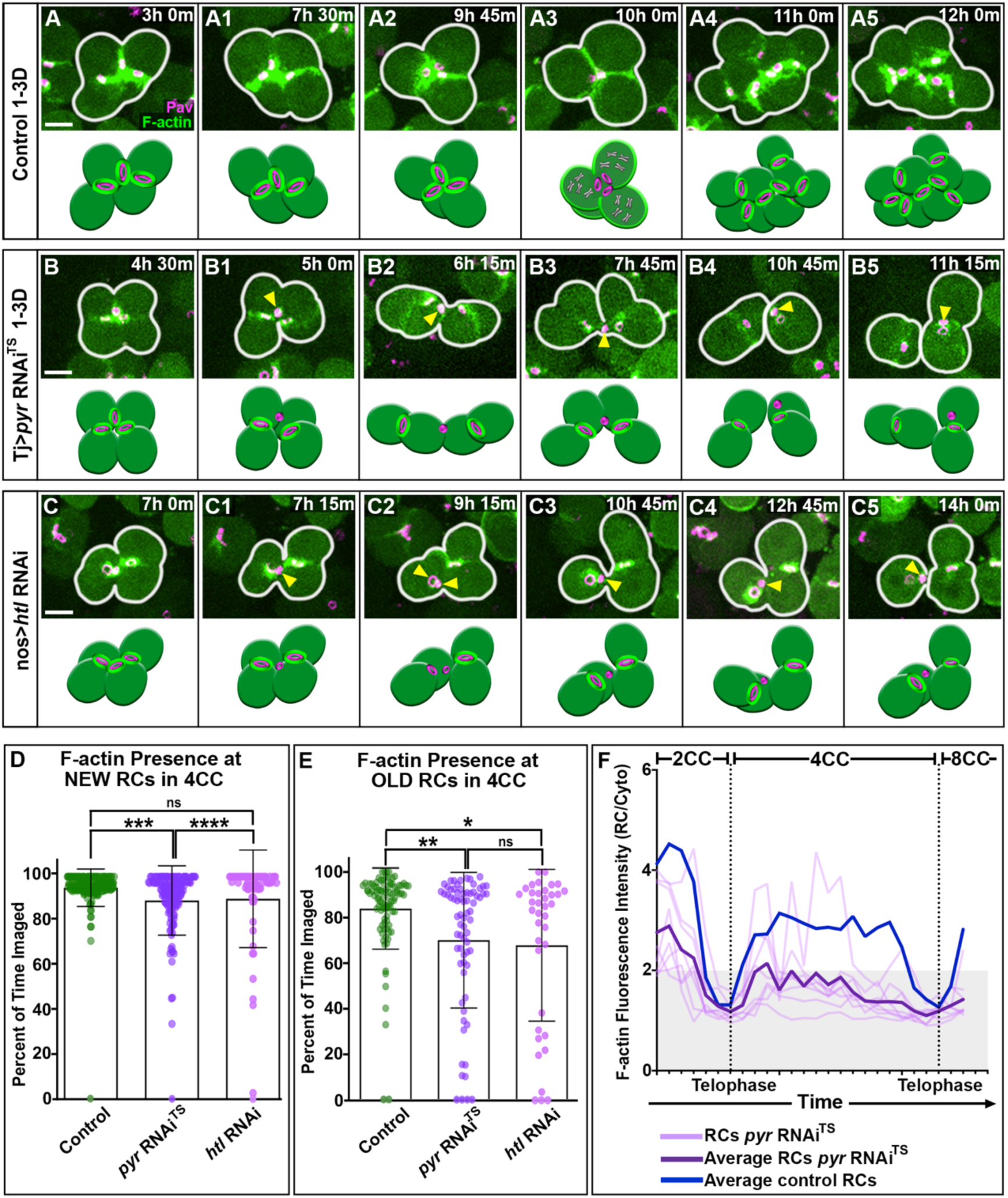
Somatic cells control F-actin stability at germline RCs via FGF signaling. (A-C) Time-lapse imaging and associated diagrams of nos-ABD-moe::GFP, Pav::mCh. Each image is 1-6 slices with images acquired every 15-minutes. (A) Control (combined 1-3D TjGal4^TS^ and nosGal4) testes exhibit stable F-actin for the duration of the 4CC cell cycle and incomplete cytokinesis. (B) 1-3D Tj>*pyr* RNAi^TS^ and (C) nos>*htl* RNAi testes both exhibit unstable F-actin at RCs (arrowhead) followed by inappropriate completion of cytokinesis. (D) Quantification of the duration of F-actin presence at new RCs over the imaging period in control (n=180), 1-3D Tj>*pyr* RNAi^TS^ (n=144), and nos>*htl* RNAi (n=76) testes. The duration of F-actin presence at control RCs is significantly different from RCs in *pyr* RNAi^TS^ (p=0.0007, ordinary ANOVA), but not from RCs in *htl* RNAi (p=0.4224, ordinary ANOVA). *pyr* RNAi^TS^ RCs have significantly reduced F-actin at new RCs compared with *htl* RNAi (p<0.0001, ordinary ANOVA). (E) Quantification of the duration of F-actin presence at old RCs over the imaging period in control (n=92), Tj>*pyr* RNAi^TS^ (n=72), and nos>*htl* RNAi (n=37). The duration of F-actin presence at old RCs in control testes is significantly different from that in *pyr* RNAi^TS^ (ordinary ANOVA, p=0.006), and *htl* RNAi (p=0.0363, ordinary ANOVA). F-actin presence at old RCs is not significantly different between *pyr* RNAi^TS^ (p=0.006, ordinary ANOVA), and *htl* RNAi (p=0.0363). (F) Quantification of normalized RC F-actin intensity over several rounds of germ cell cyst divisions in *pyr* RNAi^TS^ testes (n=8). Scale= 5 µm.

To quantify these variable defects in F-actin, we determined the duration of F-actin presence at RCs over time (Fig.3D,E). In control cysts, F-actin is retained at newly generated RCs for over 90% of the imaging period (Fig.3D). At the old RC, F-actin is maintained for ∼84% of the imaging period (Fig.3E), consistent with the slight delay in F-actin reassembly at old RCs following mitosis. Critically, there was significant reduction in duration of F-actin presence at both new and old RCs upon FGF inhibition compared with controls (Fig.3D,E). The defect was most severe at old RCs, with many maintaining F-actin for only 0-50% of the imaging period (Fig.3E).

Finally, we directly quantified F-actin fluorescence intensities at individual RCs over time in Tj>*pyr* RNAi^TS^ testes (Fig.3F). Given the severity of defects observed at old RCs, we focused our live quantification on the existing RCs of 2CCs as they transitioned from two to four to eight cells over the imaging period. Compared with old RCs from control cysts (Fig. 3F, average=blue line) in which F-actin is only absent during mitosis, we found that *pyr* depletion significantly reduced F-actin enrichment at RCs throughout the cell cycle. In fact, F-actin intensities consistently fall below the threshold indicative of F-actin absence (Fig.3F, grey rectangle). These data suggest that somatic FGF signals control germline incomplete cytokinesis by regulating maintenance of F-actin at RCs over multiple rounds of TA divisions.

### Pharmacological inhibition of FGFR kinase activity induces RC F-actin disassembly and completion of cytokinesis in germline cysts

To further interrogate the requirement for FGFR in germ cells, we used a well-established pharmacological inhibitor of FGFR tyrosine kinase activity, SU5402 (Bowles et al., 2010; Harpelunde Poulsen et al., 2019; Marques et al., 2008; Mohammadi et al., 1997). In each experiment, we imaged testes expressing: 1. no*s*-ABD-moe::GFP, Pav::mCh to visualize F-actin at RCs, and 2. nos*-*Erk Sensor, Histone::mCh to quantify FGF pathway activity over time. For all trials, testes were imaged for 3-hours prior to drug addition, at which point standard media with DMSO was removed and replaced with media containing 12.5 µM of the SU5402 FGF inhibitor. Erk activity and RC F-actin were then quantified at discrete intervals during the DMSO and SU5402 treatment periods. We observed a significant decrease in Erk activity after five hours of SU5402 treatment, which was then sustained (Fig.4A,C,E). Critically, F-actin at RCs was also dramatically reduced after SU5402 treatment, with both new and old RCs exhibiting significant F-actin depolymerization at 5 and 10 hours post drug addition (Fig.4B, D, F). Thus, loss of FGFR activity induced rapid and significant disassembly of F-actin at RCs within differentiating germline cysts.

**Fig. 4.**
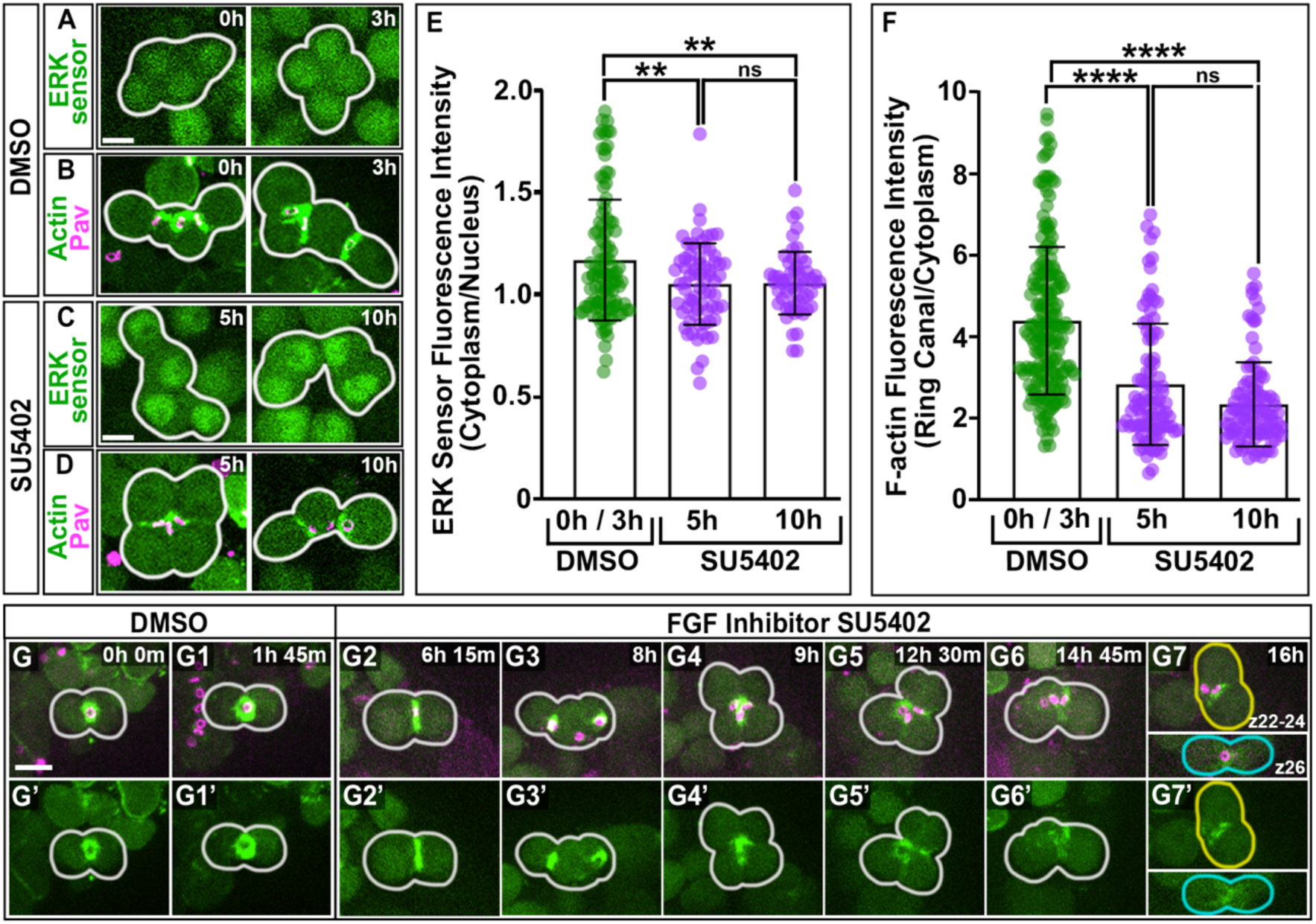
Pharmacological inhibition of FGFR kinase activity induces RC F-actin disassembly and completion of cytokinesis in germline cysts. (A-D) Time lapse imaging of 4CCs in (A) nos*-*Erk Sensor, Histone::mCh in DMSO-media at 0h and 3h. (B) nos-ABD-moe::GFP, Pav::mCh in DMSO-media at 0h and 3h. (C) nos*-*Erk Sensor, Histone::mCh in SU5402-media incubated for 5h and 10h. (D) nos-ABD-moe::GFP, Pav::mCh in SU5402-media incubated for 5h and 10h. Each image is 1-6 z-slices. Images acquired every 15 minutes. Scale=5 µm. (E) Cytoplasmic to nuclear Erk quantification in DMSO-media (n=109) and following 5h (n=71) and 10h (n=55) SU5402-media incubation. Erk activity is significantly reduced at 5h (p=0.0053) and 10h (p=0.0047, ordinary ANOVA). (F) Quantification of normalized RC F-actin intensity in DMSO-media (n=191) and following 5h (n=113) and 10h (n=109) SU5402-media incubation. RC F-actin is significantly reduced at 5h and 10h (p<0.0001, ordinary ANOVA). (G-G7”) Time-lapse imaging of one germ cell cyst through DMSO-media and SU5402-media incubation. F-actin is reduced and the cyst inappropriately completes cytokinesis (G7-G7’).

Finally, this approach permitted us to visualize changes in F-actin polymerization at RCs within the same cyst prior to drug treatment and then over time following addition of the FGFR inhibitor (Fig.4G-G7’). Similar to genetic depletion of either the Pyr ligand or Htl receptor, pharmacological inhibition of FGFR led to instability of F-actin at RCs within 4CCs over time (Fig.4G2-7’). While F-actin remains stable during the pre-drug and early post-drug imaging periods (Fig.4G-G3), both new and old RCs exhibited progressive F-actin disassembly beginning ∼5 hours after inhibition of FGFR (Fig.4G5-G6). Ultimately, the 4CC completes cytokinesis at the oldest RC, producing two independent 2CCs (Fig.4G7). Of note, while FGF inhibition dramatically impacted maintenance of RC F-actin, there was no defect in F-actin at nascent RCs nor in reassembly of F-actin at old RCs following mitotic exit. Together with our genetic analyses, these data reveal a critical role for FGF signaling in transit amplifying germ cells to promote RC stability and integrity of incomplete cytokinesis.

### Soma-germline FGF signaling is required for germline incomplete cytokinesis and regulated cyst elimination

While failure of incomplete cytokinesis is an exceptionally rare event under homeostatic conditions (4.3%, Fig5A-A4, D), we observed significant cyst abscission when FGF signaling was inhibited (Fig.3B and 3C, Fig.4G, Fig.5B,C and G). Indeed, we observed failure of incomplete cytokinesis at both new and old RCs upon somatic depletion of *pyr* (Fig.5B4 and 5C4), with nearly 16% of RCs undergoing abscission after 3 days of *pyr* depletion (Fig.5D).

**Fig. 5.**
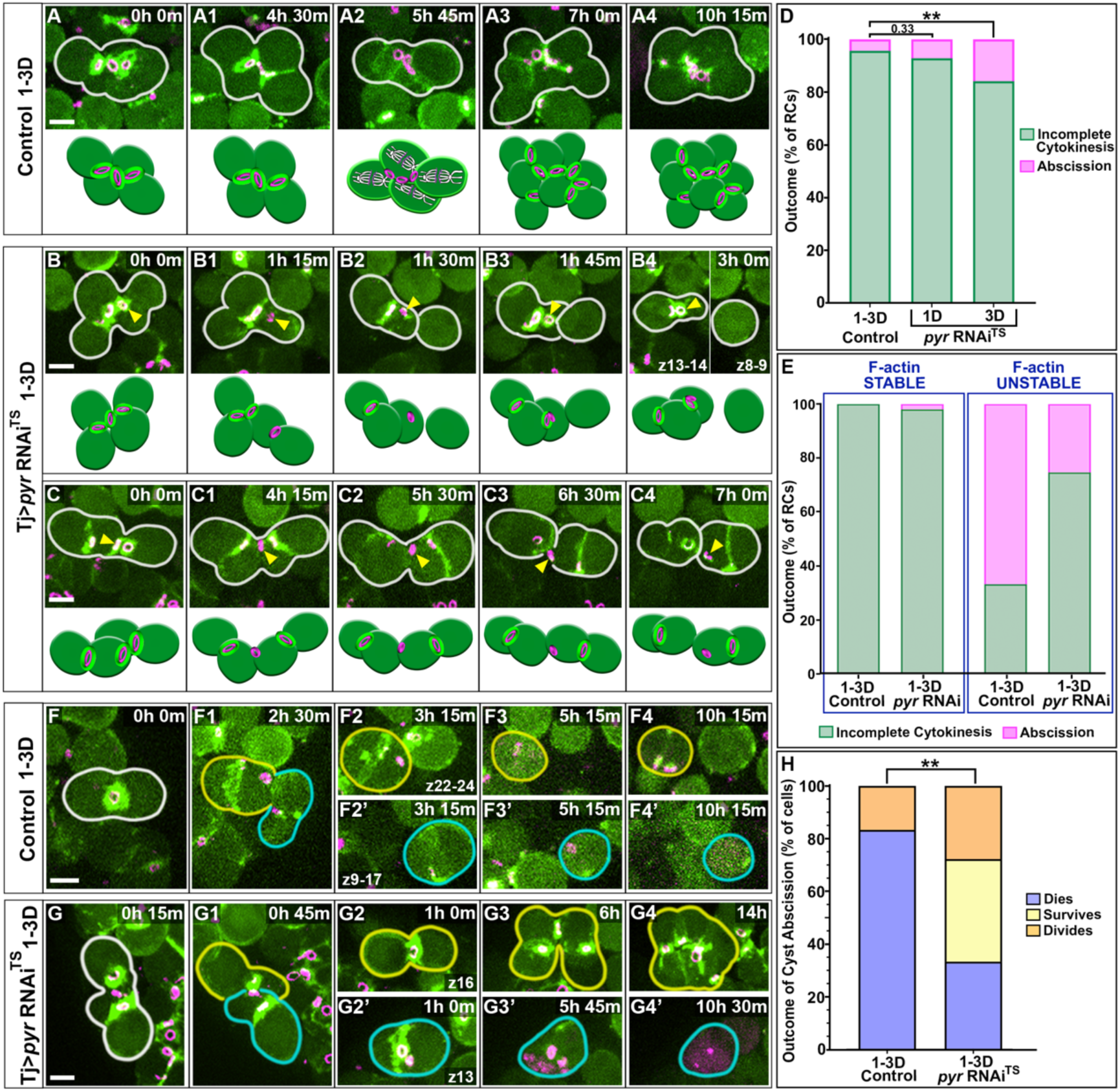
Soma-germline FGF signaling is required for germline incomplete cytokinesis and regulated cyst elimination. (A) Time-lapse imaging of 1-3D control TjGal4^TS^ 4CC-8CC division. (B) Time-lapse imaging of Tj>*pyr*RNAi^TS^ abscission at 1 new RC (arrowhead) and retention of the resulting midbody (arrowhead) (C) Time-lapse imaging of 1-3D Tj>*pyr* RNAi^TS^ abscission at oldest RC (arrowhead). (D) Quantification of the percent of RCs that execute abscission in control (n=6/138), 1D *pyr* RNAi^TS^ (n=11/153), and 3D *pyr* RNAi^TS^ (n=10/63). Abscission frequency is significantly increased with 3D of *pyr* depletion (p=0.0093), but not at 1D (p=0.33, Fisher’s Exact). (E) Quantification of the percent of RCs that execute abscission when F-actin is stable in control (n=0/93), and 1-3D Tj>*pyr* RNAi^TS^ (n=2/99), versus unstable F-actin in control (n=6/9) and 1-3D Tj>*pyr* RNAi^TS^ (n=17/75). (F) A control cyst that executes abscission followed by elimination through cell death (n=10/12). (G) 1-3D *pyr* RNAi^TS^ germ cells abscised, with one released pair undergoing apoptosis (G2’-G4’, n= 12/36) while the other cyst survived and executed mitosis (G2-G4, n=24/36). (H) Significantly fewer cysts died as the result of cyst abscission in 1-3D *pyr* RNAi^TS^ compared with controls (p=0.0059, Fisher’s exact).

To determine if cyst abscission directly correlated with unstable RC F-actin, we quantified the outcome at individual RCs over time separately for those with stable versus unstable F-actin (Fig.5E, Fig.S3). Strikingly, we found that abscission almost never occurred at RCs with stable F-actin (control, 0%; Tj>*pyr*RNAi^TS^, 2%). By contrast, cyst abscission was frequently observed at RCs with unstable F-actin in both control (66.7%) and Tj>*pyr*RNAi^TS^ testes (25.3%) (Fig.5E). This suggests that F-actin stability at RCs is essential to maintain robust incomplete cytokinesis within cysts and that loss of F-actin permits inappropriate completion of cytokinesis.

Failure of incomplete cytokinesis in germline cysts has long-term detrimental effects on both the cysts themselves and the tissue as a whole (K. L. Lu & Yamashita, 2017). Our live imaging approach allowed us to interrogate the *immediate* consequence to cysts upon completion of cytokinesis. While cyst abscission is rare in controls, when incomplete cytokinesis *does* fail nearly all released germ cells are eliminated through apoptosis (10/12, 83.3%, Fig.F-F4’,5H). Abscission at the old RC in a control 4CC resulted in release of two germ cell cysts (Fig.F2,F2’) that independently execute apoptosis, identified by morphological changes within the germ cells and then eventual breakdown of the cyst (Fig.5F2-F4’) (K. L. Lu & Yamashita, 2017; Yang & Yamashita, 2015). Thus, it seems that in the rare event of inappropriate cyst abscission during homeostasis, there is robust elimination of the resulting germ cells. Critically, we found that only 33% (12/36) of released germ cells die following an abscission event in *pyr* RNAi^TS^ testes (Fig.5G-G4’,H). The remaining 66.7% of germ cells survived for the remainder of the imaging period, and over 20% of these underwent another successful round of mitotic division (Fig.5G2-G4,H). Thus, not only does loss of FGF signaling induce significant F-actin instability and abscission at RCs, but it also prevents robust elimination of aberrantly derived cysts.

### Somatic FGF signals activate the germline-intrinsic Src64 kinase cascade to regulate RC F-actin

We next sought to identify the mechanism by which FGFR activation promotes maintenance of RC F-actin in the TA pool. Previous work has found a critical role for the cytoplasmic tyrosine kinase, Src64, in regulating RC F-actin in the *Drosophila* testis (Eikenes et al., 2015). Active Src64 phosphorylates Abelson kinase (Abl) and promotes localization of SCAR (WAVE-1) to the RC, where it then directly activates the F-actin nucleator, Arp2/3. Depletion of either kinase is sufficient to disrupt SCAR as well as F-actin at the RC (Eikenes et al., 2015). As Src-family kinases are a conserved target of active growth factor receptors, including FGFR (X. Li et al., 2004), we asked whether FGF signaling regulates RC F-actin through Src64 kinase.

To test this hypothesis, we first assayed germline kinase activity by visualizing phosphorylated-Tyrosine (pTyrosine). Previous work found that RCs accumulate increasing amounts of tyrosine phosphorylated proteins during TA divisions and that this pTyrosine enrichment is dependent upon Src64 kinase activity (Eikenes et al., 2015). We found that pTyrosine is indeed enriched at old RCs in control germline cysts (Fig.6A-6A”) (Eikenes et al., 2015; Gary R. Hime et al., 1996; Robinson & Cooley, 1996). Excitingly, we find that pTyrosine epitopes are significantly reduced at all RCs following just 1-3 days of somatic *pyr* depletion (Fig.6B-6B”,C). Thus, loss of the FGF ligand in somatic cells is sufficient to reduce overall germline tyrosine kinase activity.

**Fig. 6.**
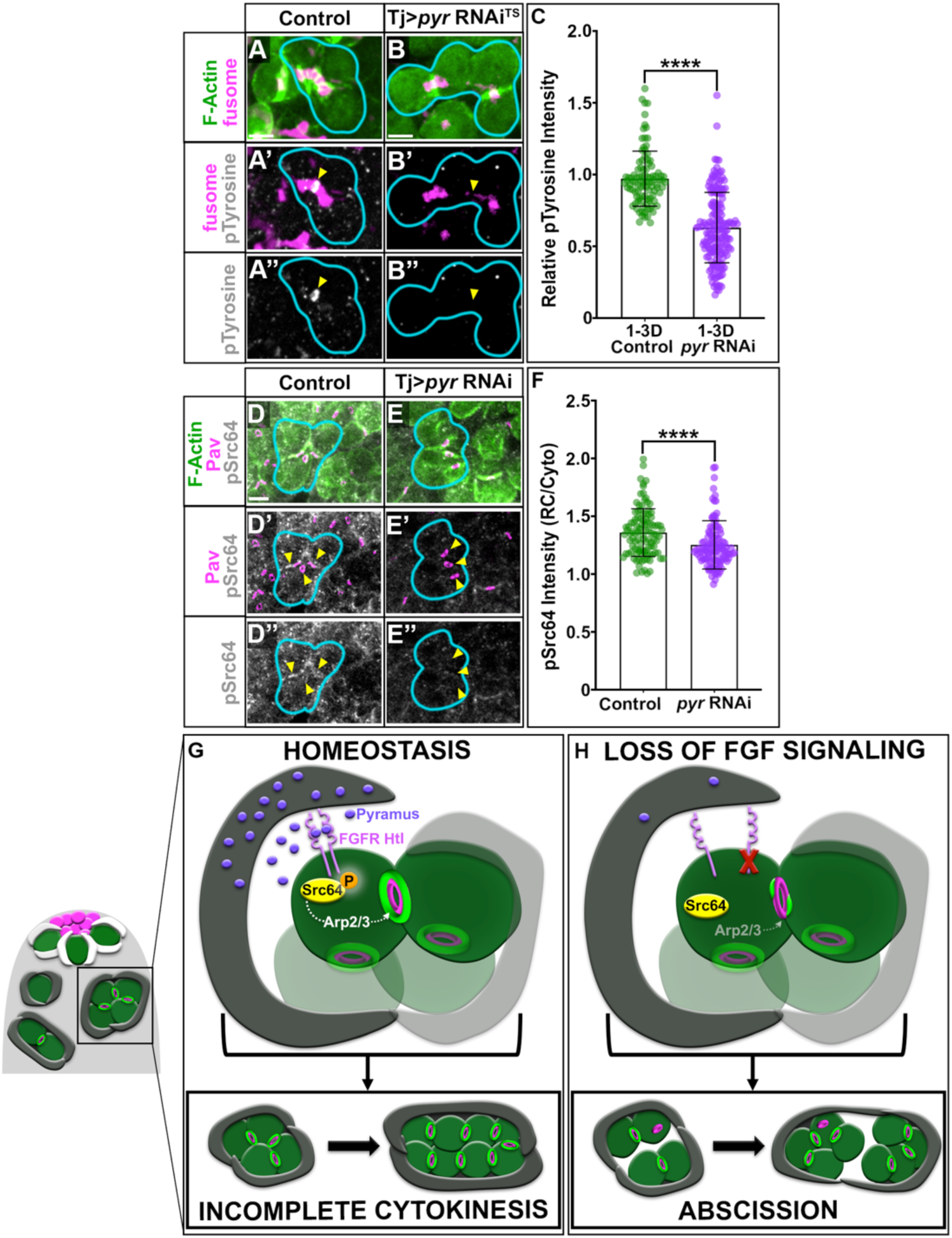
Somatic FGF signals activate the germline-intrinsic Src64 kinase cascade to regulate RC F-actin. (A) pTyrosine stain in control 1-3D TjGal4^TS^ and (B) 1-3D Tj>*pyr* RNAi^TS^ cysts. F-actin (green), 1b1 (magenta), and pTyrosine (grey). Scale=5µm. Arrowheads indicate location of old RC. (C) Relative pTyrosine intensity at control (n=101) and Tj>*pyr* RNAi^TS^ RCs (n=167) (p<0.0001, Mann-Whitney t-test). (D) pSrc64 stain in control TjGal4 and (E) Tj>*pyr* RNAi cysts. F-actin (green), Pav (magenta), and pSrc64 (grey). Scale=5µm. Arrowheads indicate all RCs. (F) Normalized pSrc64 intensity at RCs in control (n=106) and Tj>*pyr* RNAi RCs (n=111) (p<0.0001, Mann-Whitney t-test).

To specifically determine if FGFR activates Src64 in the testis, we assessed levels of phosphorylated Src64 protein by immunofluorescence. As expected, we found that pSrc64 accumulated at all RCs within germline cysts (Fig.6D-6D”, arrowheads). Strikingly, inhibition of *pyr* in somatic cells significantly reduced the amount of pSrc64 localized to 4CC RCs (Fig.6E-E’’, arrowheads; Fig.6F). These data suggest that somatic FGF signaling controls germline RC stability via activation of the germ cell intrinsic kinase cascade required for maintenance of F-actin integrity at RCs (Fig.6G,H).

## Discussion

Transit-amplification combined with incomplete cytokinesis is a deeply conserved feature of germ cell biology from *Hydra* to humans (Haglund et al., 2011; Price et al., 2023). Historically, studies of incomplete cytokinesis have lacked the spatio-temporal resolution to address how this process occurs reiteratively over multiple rounds of TA divisions. Our live imaging now reveals this critical process in the differentiating germline is both dynamically regulated and directly controlled by somatic support cells. By imaging germline cysts through multiple mitotic divisions, we discovered distinct F-actin dynamics during incomplete cytokinesis at new versus old RCs. We confirmed the CR origin of F-actin at new RCs and identified a previously unappreciated, *de novo* F-actin polymerization event required at old RCs following mitosis. Critically, we find that maintenance of F-actin at all RCs is regulated non-autonomously via FGF signaling from associated somatic cells and that FGFR activation initiates the Src64-Abl-Arp2/3 cascade required for F-actin stability at the ICB (Fig.6G). Finally, we show that unstable RC F-actin leads to failure of incomplete cytokinesis within the germline and execution of abscission within cysts (Fig.6H). Together, these data suggest a crucial role for conserved growth factor signaling and somatic support cells in maintaining germline RC F-actin and ultimately, integrity of the transit amplifying pool.

### Somatic support cells control germline incomplete cytokinesis through FGF signaling and stabilization of F-actin at RCs

Execution of incomplete cytokinesis by the germline and formation of stable RCs has historically been viewed as an arrest of traditional cytokinesis. In accordance with recent work (Price et al., 2023), this study reveals that RC stability across TA divisions is controlled by a regulatory program initiated by somatic support cells and distinct from canonical cytokinesis. Our live imaging of germline cysts across multiple mitotic divisions revealed two distinct F-actin polymerization events at RCs – stabilization of CR F-actin at new RCs and *de novo* F-actin polymerization following mitotic exit at old RCs. Despite these differences in origin, FGF is required for persistent F-actin maintenance at both new and old RCs. Intriguingly, however, the F-actin defects induced by loss of FGFR activity exclusively impact F-actin *maintenance* but not *initial polymerization*. With all genetic and pharmacological inhibitions of FGF signaling, we still observed robust polymerization of F-actin at new RCs *and* reestablishment of F-actin at old RCs in nearly 100% of cysts. Yet, ∼40% of all RCs exhibited defects in F-actin maintenance over time (Fig.S3). These data strongly suggest that initial establishment of F-actin at the RC is FGF and Src64 independent. Instead, F-actin assembly at RCs is likely mediated through canonical cytokinetic effectors such as RhoA and formins (Glotzer, 2003; Gunsalus et al., 1995; Pollard, 2010). By contrast, *maintenance* of RC F-actin is FGF-dependent through the effector Src64 kinase and, ultimately, Arp2/3 (Fig.6G,H).

Thus, much like the midbody remodeling that occurs during RC formation (Price et al., 2023), F-actin regulation combines canonical cytokinetic factors with unique regulatory components to ensure robust execution of stable incomplete cytokinesis.

Importantly, while loss of FGF signaling induces F-actin instability and abscission at both new and old RCs, we consistently observed more penetrant defects at the old RC. Indeed, nearly 70% of all abscission events induced by loss of FGF activity occurred at the old RC. This increased severity likely reflects the novel origin and dynamic nature of F-actin at old RCs, as this F-actin pool must be reestablished after every TA division. Initial stabilization of old RC F-actin may require additional crosslinking factors that take time to accumulate after mitotic exit, making this F-actin pool more sensitive to loss of FGF signals and decreased Src64 activity.

### A conserved role for stable F-actin in preventing completion of cytokinesis

Previous work identified a requirement for RC F-actin in maintenance of proper RC diameter (Eikenes et al., 2015). However, these studies were conducted exclusively in fixed tissue and thus could not address the consequence of F-actin instability at RCs over time. Our longitudinal imaging permitted us to directly examine the consequence of unstable RC F-actin to germline cysts. This approach revealed a direct link between F-actin maintenance at RCs and integrity of incomplete cytokinesis. Loss of somatic FGF signaling caused dramatic but variable defects in RC F-actin maintenance; in 2- and 4-cell cysts, about half of all RCs retained relatively stable F-actin while the other half exhibited severe F-actin defects (Fig.3B-F). Critically, while nearly all RCs with stable F-actin properly execute incomplete cytokinesis, about 20% of RCs with unstable F-actin inappropriately undergo abscission (Fig.5E). This strongly suggests that stable and persistent F-actin at RCs is essential for proper execution of the incomplete cytokinetic program and prevention of cyst abscission.

In animal cells from *Drosophila* to humans, F-actin must be cleared from the ICB during late cytokinesis for abscission to complete (Abe et al., 1996; Gunsalus et al., 1995; Kaji et al., 2003; Nagaoka et al., 1996; Pollard, 2010; Somma et al., 2002). Indeed, cells utilize polymerization of F-actin at the ICB as a mechanism to *prevent* cytokinesis progression. Across all cell types, inefficient chromosome segregation during mitosis induces the “No-Cut” pathway which blocks abscission in part through polymerization of F-actin at the cytokinetic ICB (Steigemann & Gerlich, 2009). In addition, our lab has shown that F-actin clearance from the germline stem cell ICB is critical for cytokinesis completion. Recruitment of abscission machinery occurs only *after* F-actin is disassembled, and retention of F-actin at the GSC ICB results in abscission failure (Lenhart & DiNardo, 2015). Thus, persistent F-actin at ICBs is a conserved mechanism to prevent abscission in both canonical and incomplete cytokinesis programs. How F-actin at RCs prevents recruitment and function of abscission machinery remains unknown. However, recent work has identified an essential role for the deubiquitinase Usp8 in execution of incomplete cytokinesis by targeting ESCRT III component Shrub to prevent abscission at RCs (Mathieu et al., 2022). In the future, it will be interesting to examine if F-actin at RCs is required for recruitment of Usp8 to the ICB and if the recruitment of abscission machinery observed upon Usp8 inhibition requires a coincident loss of F-actin at RCs.

### Failure of incomplete cytokinesis and consequences to tissue homeostasis

The highly conserved nature of incomplete cytokinesis across animal species suggests an intimate link between germline interconnectivity and effective reproduction. In the female fly, there is an explicit requirement for cytoplasmic sharing to produce a functional oocyte. In the male, cytoplasmic sharing promotes robust detection and elimination of genomically damaged germ cells by inducing apoptosis in all cells within a cyst (K. L. Lu & Yamashita, 2017). This “all or none” response is proposed to ensure robust genomic integrity of the germline by eliminating any cells with potential DNA damage, even if that damage would typically be insufficient to induce apoptotic pathways. Experimentally decreasing germline interconnectivity reduced detection and elimination of damaged germ cells (K. L. Lu & Yamashita, 2017). Thus, failure of incomplete cytokinesis and abscission of germline cysts directly impacts the ability of damaged germ cells to be culled from the TA pool. In the future, it will be important to determine if FGF inhibition yields gametes with genomic defects, or if loss of FGF and increased abscission creates a sensitized background in which tumors more readily develop with age.

The “all or none” apoptotic response to DNA damage may also function to cull germ cells that undergo inappropriate abscission events. We find that while fragmentation events are rare in control testes, individual pairs produced from complete cytokinesis are almost always eliminated through cell death (Fig.5H), retaining all-or-none elimination. By contrast, germ cells that fractionate in *pyr* depleted testes exhibited heterogenous outcomes and, importantly, only ∼30% of abscission events lead to death of the resulting separated cysts (Fig.5H). The remaining 70% of cysts survive and often perform another round of transit amplification (Fig.5H).

There are multiple reasons why continued survival of cysts post-abscission could negatively impact the tissue. First, it remains unclear whether fractionating germ cells retain their “mitotic count”. In mammals and *Drosophila*, precise control over transit amplifying mitoses is critical for homeostasis and spermatogenesis. Dysregulation in the transit amplifying pool can promote tumorigenesis or tissue atrophy (Amy A. Kiger et al., 2000; Bowles et al., 2010; Herrera & Bach, 2018; Insco et al., 2009; Reya et al., 2001; Schwitalla et al., 2013). Whether cyst abscission disrupts mitotic count is an outstanding question for future study.

Second, we found that cyst abscission routinely causes internalization and retention of the RC midbody remnant in one germ cell of the separated cysts. At the niche, GSC abscission leads to release of the cytokinetic midbody into the extracellular space (Lenhart & DiNardo, 2015), where it is then phagocytosed and degraded by adjacent somatic cells (Salzmann et al., 2014). Thus, during homeostasis, germ cells never inherit the midbody. Critically, upon loss of FGF signaling in the testis, we find that 0% of cysts release the midbody following abscission. Instead, one germ cell pair *always* retains the internalized midbody remnant, which is never degraded and is detectable hours after germ cell abscission (Fig.3 and Fig.5, arrowheads). The midbody contains hundreds of proteins, and inheritance of the organelle has been shown to regulate cell identify, differentiation, proliferation and tumorigenesis (Dionne et al., 2015; Farmer, 2022; H. Li et al., 2026). How inappropriate retention of the midbody might modify or disrupt differentiating germ cell biology remains entirely unknown. In addition, our lab has observed that retention of internal midbodies (in GSCs and gonia) can disrupt subsequent mitoses, with germ cells undergoing multipolar divisions (Lenhart lab unpublished; Fig.S4). This suggests that internal midbodies may become activated as microtubule organizing centers; an intriguing possibility considering the significant overlap in protein composition at the midbody and activated centrosome (Fabbro et al., 2005; Martinez-Garay et al., 2006; Morita et al., 2010). Supernumerary centrosomes are also often observed in cancer cells and can induce multipolar spindle formation and aneuploidy (Kalkan et al., 2024; Shin et al., 2021). Thus, in the future, it will be critical to explore the impact of internalized midbodies on germline cell biology and homeostasis. **Somatic control of germline incomplete cytokinesis may be conserved**

It is well established that somatic support cells regulate multiple aspects of germline biology across species (Amy A. Kiger et al., 2000; Fabrizio et al., 2003; Fairchild et al., 2015; John Tran et al., 2000; Lenhart & DiNardo, 2015; Masaki et al., 2018; Oatley & Brinster, 2012; Takashima et al., 2015; Tang et al., 2017). Our work suggests somatic FGF signaling may be another potentially conserved aspect of gametogenesis. In the mammalian testis, at least two FGF ligands are secreted by somatic Sertoli cells (FGF2 and FGF9) (Bowles et al., 2010; Masaki et al., 2018; Takashima et al., 2015). Both ligands have been shown to regulate germline stem cells as well as mitotically dividing spermatogonia (Bowles et al., 2010; Masaki et al., 2018; Takashima et al., 2015). As live imaging of the mouse testis becomes technically more accessible, it will be important to examine if somatic FGF signals are also required for long-term maintenance of RC stability during TA divisions in mammals. Indeed, the FGFR effector Src64 in the fly shares significant functional similarities with the mammalian tyrosine kinase TEX14 (Cooley, 1998; Greenbaum et al., 2006; Iwamori et al., 2010; N. Lu et al., 2004). As no upstream regulators of TEX14 have been identified, future work interrogating the degree of conservation between the fly and mouse incomplete cytokinetic programs should directly assess if TEX14 might be a target of Sertoli cell-derived FGF signaling.

## Methods

| REAGENT or RESOURCE | SOURCE | IDENTIFIER |
| --- | --- | --- |
| **Antibodies** |  |  |
| Chicken anti-GFP | Aves Labs | RRID: AB10000240 |
| Mouse anti-1B1 | DSHB | RRID: AB528070 |
| Chicken anti-Pyramus<sup>intra</sup> | Angeline Stathopoulos | NA |
| Rabbit anti-pSrc64 | Alana M O' Reily | NA |
| Rabbit anti-Vasa | Boster Biological Technology Co. | RRID: DZ 41154 |
| Rat anti-DE-cadherin | DSHB | RRID: AB 528120 |
| Rabbit anti-pTyrosine | Sigma-Aldrich | T1325 |
| **Chemicals, peptides, and recombinant proteins** |  |  |
| Phosphatase inhibitor cocktail 2 | Sigma-Aldrich | P5726 |
| Su5402 FGF Inhibitor | Sigma-Aldrich | 572630 |
| **Experimental models: Organisms/strains** |  |  |
| *D. melanogaster*: Pav::mCh | Cooley Lab | NA (doi-10.1016) |
| *D. melanogaster*: Tj-Gal4 | Kyoto Stock Center | KSC: 104055<br>FBti0034540 |
| *D. melanogaster*: nos-Gal4-VP16 | Gift from Ruth Lehmann and Erica Selva | N/A |
| *D. melanogaster*: nos-ABDmoe::GFP | The Lehmann Lab | N/A |
| *D. melanogaster*: UAS-CD8::GFP | Bloomington *Drosophila* Stock Center | BDSC: 32186<br>FBti0131963 |
| *D. melanogaster*: UAS-pyr RNAi | Bloomington *Drosophila* Stock Center | BDSC: 63547<br>FBgn0033649 |
| *D. melanogaster*: Erk-KTR::Clover, His::mCh | Bloomington *Drosophila* Stock Center | BDSC: 95287<br>FBti0226349 |
| *D. melanogaster*: UAS-Red Stinger | Bloomington *Drosophila* Stock Center | BDSC: 8546<br>FBti0040829 |
| *D. melanogaster*: Tj-Gal4, Tub-Gal80<sup>TS</sup> | Recombinant made from Tj-Gal4 and Gal80<sup>TS</sup> (Erika Bach) | N/A |
| *D. melanogaster*: Pyr-Gal4 | Bloomington *Drosophila* Stock Center | BDSC: 91292<br>FBti0212964 |
| *D. melanogaster*: UAS-*htl* RNAi | Bloomington *Drosophila* Stock Center | BDSC: 57313<br>FBgn0010389 |
| **Software and algorithms** |  |  |
| ImageJ/Fiji | https://fiji.sc | RRID: CVCL_0299 |
| Graphpad Prism | https://www.graphpad.com/scientific-software/prism/ | RRID:SCR_002798 |
| Adobe Photoshop | https://www.adobe.com/uk/products/photoshop.html | RRID:SCR_014199 |
| **Other** |  |  |
| Imaging dishes with coverslip bottom | MatTek | Cat# P35G-1.5-14-C |
| Olympus iX83 inverted spinning disk confocal | Olympus/Evident | N/A |
| Hamamatsu EM-CCD camera | Olympus/Evident | N/A |
| 60X 1.4NA silicon oil immersion objective | Olympus/Evident | N/A |

### Fly Stocks and Crosses

*Drosophila* stocks were maintained on Bloomington Drosophila Stock Center (BDSC) standard cornmeal medium in vials or bottles. All crosses without a temperature-sensitive GAL80 were raised at 25°C and 1–3-day old adult males were selected for experimentation. Temperature-sensitive GAL80 flies were grown at 18°C, selected for 1-3 days, and transferred to 29°C for 1 or 3 days to induce RNAi-mediated knockdown. For all experiments, controls are age-matched and temperature-matched sibling and parental flies.

### Time Lapse Imaging

Extended time-lapse imaging and culture conditions were performed as previously described (Lenhart et al., 2019; Lenhart & DiNardo, 2015; Rebecca Sheng & Matunis, 2011; Roach et al., 2025; Roach & Lenhart, 2024). Age and temperature-matched samples were dissected in Ringer’s solution and mounted onto a poly-lysine coated coverslip (imaging dishes, MatTek). Ringer’s solution was removed and imaging media (15% fetal bovine serum, 0.5x penicillin/streptomycin, 0.2 mg/ml insulin in Schneider’s insect media) was added. Samples were imaged every 5 or 15-minutes for up to 24-hours on an Olympus iX83 with a Yokagowa CSU-10 spinning disk scan head, 60x 1.4 NA silicon oil immersion objective and Hamamatsu EM-CCD camera using 1 µm z-step size (40 µm stacks). Experiments were repeated a minimum of two times and at least eight samples were analyzed for each genotype/condition. Time-lapse images were analyzed and z-projections generated using Fiji software.

### Analysis of RC F-actin from Live Imaging

#### Quantitative F-actin Measurement (RC / Cytoplasm)

To investigate RC F-actin dynamics across reiterative mitoses, we measured the mean intensity of RC F-actin across the 2CC-8CC stages. At each time point, RC F-actin was normalized to the mean intensity of cytoplasmic F-actin intensity within germ cells of the same cyst, combined with background subtraction. We then aligned each cyst by their mitotic telophase values in order to observe population-level trends in the F-actin cytoskeleton across germ cell mitoses.

#### Leveraging GSC cytokinesis to determine F-actin presence versus absence threshold

In order to quantitatively categorize intensity measurements as “F-actin present” vs “F-actin absent” at RCs, we leveraged completely dividing germline stem cells (GSCs). Our lab has previously shown that GSCs execute a modified but complete cytokinetic program (Lenhart & DiNardo, 2015). We quantified the intensity of nos-ABD-moe::GFP at the ICB relative to cytoplasmic signal from the same GSC-daughter pair combined with background subtraction. We then quantified this ratio over time, in order to experimentally determine the threshold amount of F-actin localized to the ICB prior to and during abscission. By calculating the 95% confidence intervals for GSC F-actin intensity across cytokinesis, we determined that F-actin can be assigned “present” at the RC if the RC/Cyto ratio is greater than 2.0. F-actin ratios below 2.0 indicate F-actin disassembly associated with cytokinesis progression (Fig.S1A-S1B). These quantifications were made in both live and fixed samples, yielding identical confidence intervals with a consistent 2.0 ratio cutoff for F-actin presence at the ICB.

#### Assessment of percent of time F-Actin is Present / total time observed

To experimentally determine the duration of time F-actin persists at the RC across the total observation time, we performed qualitative assessment of RC F-actin across divisions. All 4CCs included in the analyses were observed for 4 hours or longer. RC F-actin was qualitatively assessed as present or absent by the lack of nos-ABD-moe::GFP enrichment at the Pav::mCh labelled ring canal (Fig.3B3, arrowhead). The total time F-actin was recorded as absent was then subtracted from the total observation time, and from this, “Percent Time F-actin Presence” was calculated for each RC.

#### Qualitative Binary Stable / Unstable Analysis

To assess abscission frequency as a function of F-actin stability, we categorized F-actin enrichment at each RC tracked across the 4CC cell cycle. RCs were assessed as retaining stable F-actin if nos-ABD-moe::GFP remained polymerized at the Pav::mCh ring for the duration of interphase. RCs were categorized as exhibiting unstable F-actin if nos-ABD-moe::GFP was visibly absent at Pav::mCh rings for a minimum of 30-minutes during interphase. We restricted this analysis to 4CCs that were imaged for a minimum of 4-hours.

### Analysis of RC F-actin from Fixed Imaging

#### Quantitative F-actin Measurement (RC / Cytoplasm)

Just as with live samples, we quantified the fluorescence intensity of nos-ABD-moe::GFP at RCs relative to the intensity within cytoplasm of the same cyst combined with background subtraction. All RCs in 2CC-4CC stage gonia were included in quantification.

#### Analysis of Abscission from Live Imaging

Abscission events were quantified from live imaging through observation of RC condensation, followed by movement of the midbody remnant away from the previously existing ICB. Separation of cysts was confirmed by physical separation of the two released germ cell clusters and loss of all detectable cytoplasmic and membrane connections. A similar methodology was utilized to study developmentally-regulated cyst fractionation in the mouse testis (Lei & Spradling, 2013).

### Immunostaining

Immunostaining was performed as previously described (Lenhart et al., 2019; Lenhart & DiNardo, 2015; Roach et al., 2025; Roach & Lenhart, 2024). Briefly, testes were dissected in Ringers solution and fixed for 30 minutes in 4% formaldehyde in Buffer B (75 mM KCl; 25 mM NaCl; 3.3 mM MgCl_2_; 16.7 mM KPO_4_) followed by several washes in PBSTx (1× PBS, 0.1% Triton-X 100) and blocking in 2% normal donkey serum.

Testes were incubated in primary antibodies at 4°C at least overnight, washed several times, and then incubated in appropriate secondary antibodies for 1-hour at room temperature. After additional washes, testes were equilibrated in a solution of 50% glycerol and then mounted on slides in a solution of 80% glycerol. Primary antibodies used include: mouse anti-1B1 (DSHB, 1:50), chick anti-GFP (Aves Labs, 1:1000), chick anti-Pyramus^intra^ (Stathopoulos Lab, 1:300), rabbit anti-pSrc64 (Alana M. O’Reily, 1:100), rabbit anti-Vasa (Sigma-Aldrich, 1:5000), rat anti-DE-cadherin (DSHB, 1:20), rabbit anti-pTyrosine (Sigma-Aldrich, 1:300). Secondary antibodies used were from Jackson ImmunoResearch and used at 1:125 dilution: Alexa fluor-488 (anti-chicken 703-545-155), -Cy3 (anti-mouse 715-165-151), and -Cy5 (anti-rat 712-605-153 and anti-rabbit 711-175-152). All antibodies have been previously verified by the *Drosophila* community.

Immunolabelling against pSrc64 was performed as described above but dissections were conducted on ice and in solutions containing a phosphatase inhibitor cocktail (Sigma-Aldrich, P5726, 1:100) and each incubation/wash step was carried out at 4°C.

### Quantification of Fluorescence Intensities

#### pSrc64

Regions of interest were drawn around the broadest plane of the ring canal, and the mean fluorescence intensity was measured in Fiji. All RC measurements were normalized to the mean cytoplasmic fluorescence intensity of the cyst.

#### Pyramus^intra^

Mean fluorescence intensity was measured at the testis anterior (Fig.2A-E) in Image J. The region of interest (yellow outline) included the stem cell niche and all mitotic gonia, identified by dense Vasa+ nuclei. Final Pyr intensity = Pyr^intra^ mean intensity – background Pyr^intra^ mean intensity.

#### pTyrosine antibody measurements

Using Fiji, regions of interest were drawn around the broadest plane of the ring canal. To account for between tissue variability in immunofluorescent staining efficiency, the mean fluorescence intensity of pTyrosine at each RC was normalized to the average intensity of pTyrosine at RCs of the 16CC stage. In the absence of prominent pTyrosine staining in *pyr* RNAi^TS^ testes, the region of interest was determined using fusome and cyst morphology.

#### nos-Erk-KTR::Clover, His::mCh Biosensor

Regions of interest were drawn using the His::mCh signal to identify nuclei of each cell. The integrated density of nuclear Erk::mClover was quantified and cytoplasmic intensity was calculated by subtraction of Erk::mClover intensity within the nucleus from the whole-cell integrated density followed by background subtraction. Quantifications are reported as the cytoplasmic to nuclear ratio.

### Statistical analysis and image processing

Time-lapse images were analyzed and z-projections generated using ImageJ software. All graphical representations of data and statistical analysis were performed in Graphpad Prism (non-parametric Mann-Whitney t-test, or ordinary ANOVA). Outlier analysis (ROUT Q=0.1%) was performed for all data sets except for Fig.3D-E. Error bars represent standard deviation. *n* and *P* values are indicated in figure legends. All assays include a minimum of 2 trials and 10-testes per condition. Figures were generated using BioRender.com and Adobe Photoshop.

## Acknowledgements

We would like to thank the Lehmann, Cooley, Stathopoulos, Bach and O’Reilley labs for sharing *Drosophila* stocks and reagents. A special thanks to Dr. Stephen DiNardo for his time and expertise isolating FGF and control transcripts from ss-RNA sequencing (Raz et al., 2023). We also thank Kathy Lindsley from Evident Scientific and Drexel’s Cell Imaging Center for their technical support with confocal imaging. This work was supported by the National Institutes of Health (R01 GM138705 to K.F.L) and the Hevolution Foundation (HF-GRO-23-1199154-38 to K.F.L.).

