## Supplemental Figures for "Somatic cells non-autonomously control germline incomplete cytokinesis through FGF signaling"

**Running Title: FGF regulates germline F-actin**


Beth Kern, Zachary Y. Berkley, Samuel Price, Kari F. Lenhart

Biology Department, Drexel University


### SUPPLEMENTAL FIGURES

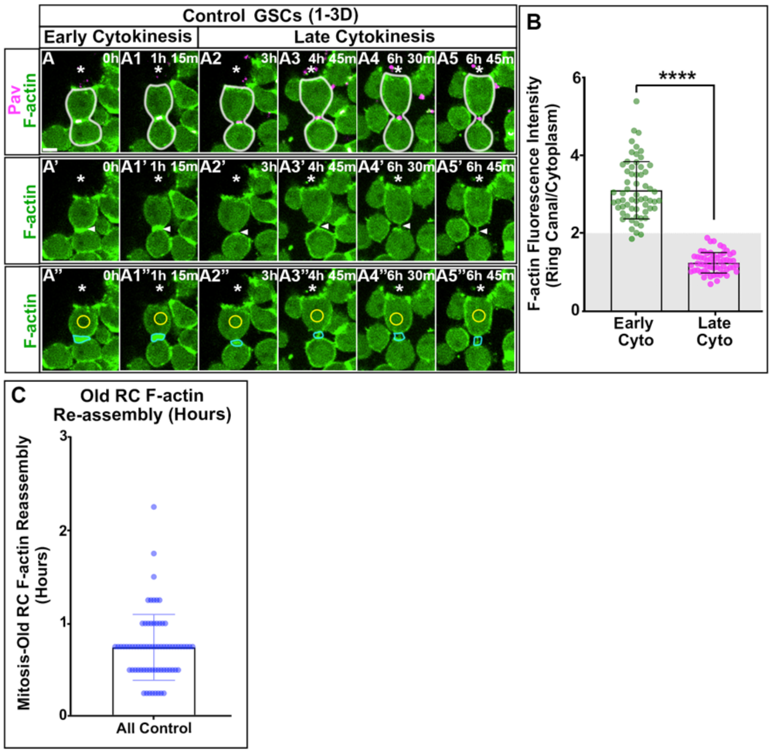


**Fig.S1 RC F-actin is highly regulated over the cyst cell cycle.** (A)Time-lapse imaging of nos-ABD-moe::GFP, Pav::mCh control GSC complete cytokinesis. (B) F-actin quantified during early cytokinesis (n=61) and late cytokinesis (n=58) ($p<0.0001$, Mann-Whitney t-test). (C) Duration of time between mitosis and old RC F-actin re-assembly in all controls (n=71). Scale= 5 µm.

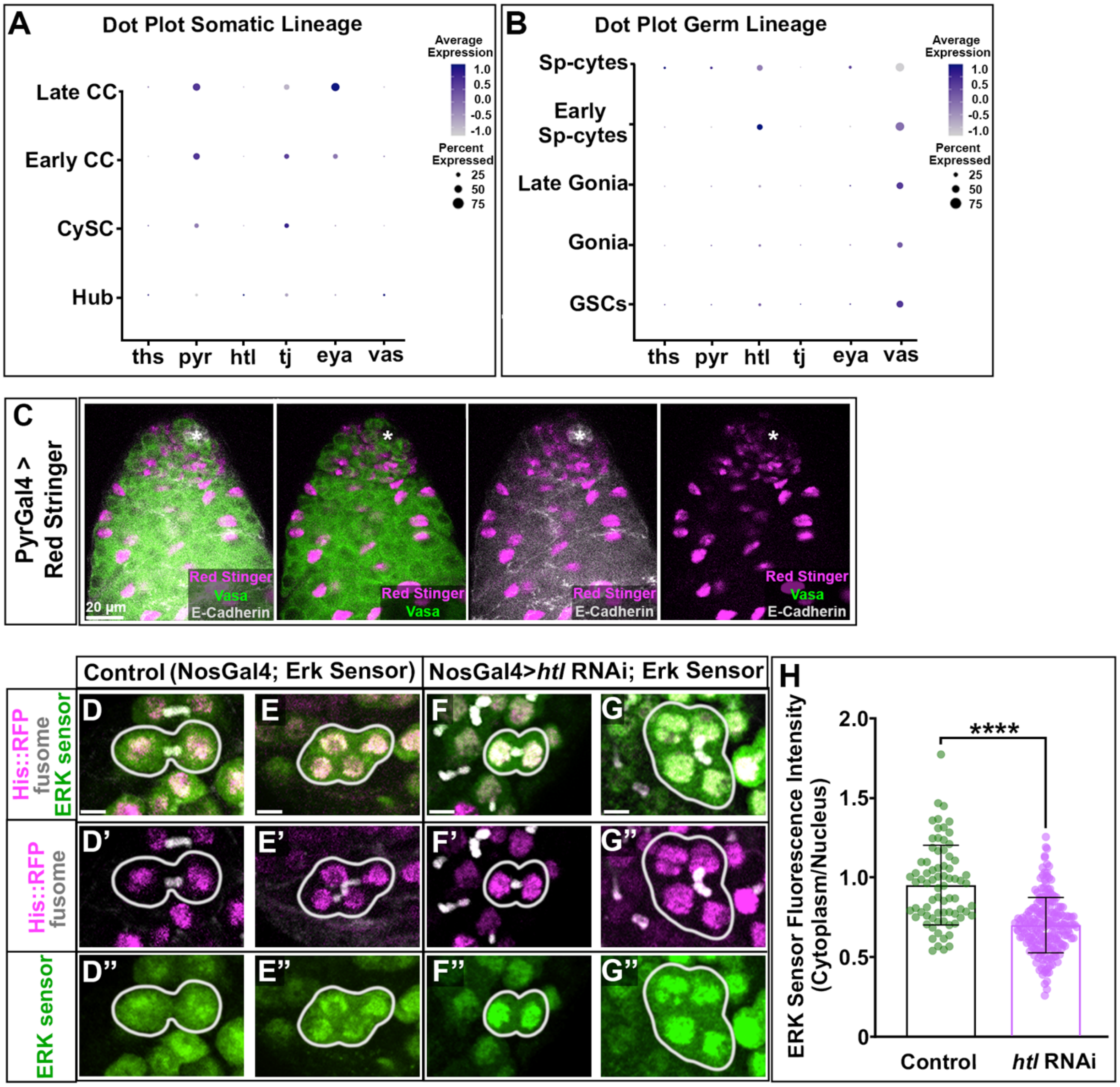


**Fig.S2. FGF ligand Pyramus is expressed by somatic cells and activates FGFR in the germline.** (A) Dot plot isolated from ssRNA sequencing of the somatic lineage (Raz et al., 2023) for *thisbe (ths), pyramus (pyr),* and *heartless (htl)* as well as control transcripts *traffic jam (tj), eyes absent (eya),* and *vasa (vas)*. (B) Dot plot of germline lineage (Raz et al., 2023). (C) PyrGal4>Red Stinger testis. Red Stinger (magenta), Vasa (green), E-cadherin (grey). (D-G" Erk-KTR::Clover, His::mCh with fusome in grey. 2CC and 4CC in (D-D", E-E") control *nos*-Gal4 and (F-F", G-G") *nos*>*htl* RNAi testes. Scale= 5 µm. (H) Cytoplasmic to nuclear Erk quantification control (n=75) and *htl* RNAi testes (n=204) (p<0.0001 Mann Whitney t-test).

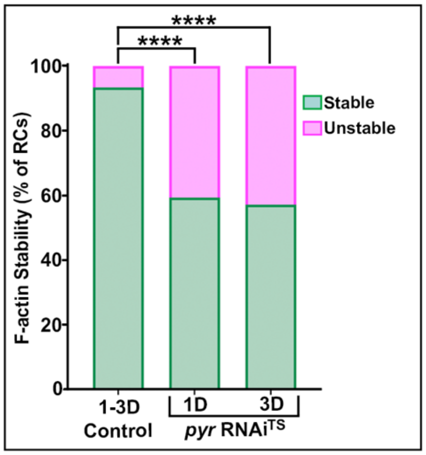


**Fig.S3. FGFR is required for germline incomplete cytokinesis and regulated cyst elimination.** (A) Quantification of the percent of RCs that exhibit F-actin instability in control (n=8/121), 1D *pyr* RNAi$^{TS}$ (n=50/123), and 3D *pyr* RNAi$^{TS}$ (n=48/112). F-actin instability is significantly increased with 1D (p<0.0001) and 3D (p<0.0001, Fisher's Exact) of *pyr* depletion.

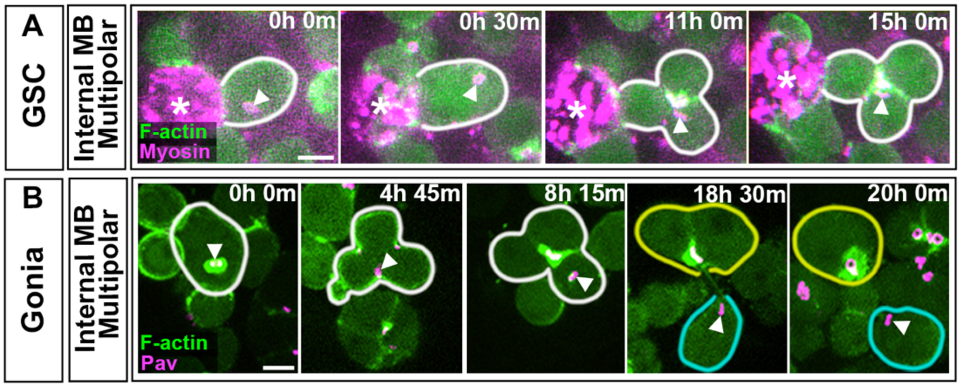


**Fig.S4. Internal midbodies can result in multipolar divisions in GSCs and gonia.** (A) Time-lapse imaging of nos-ABD-moe::GFP, Myosin::mCherry. GSC midbody internalization and subsequent multipolar division in a 21D aged testis. Scale= 5 µm. (B) Time-lapse imaging of nos-ABD-moe::GFP, Pav::mCh in a nos>*htl* RNAi testis. Gonial cell with an internalized midbody and subsequent multipolar division followed by cyst abscission. Scale= 5 µm.

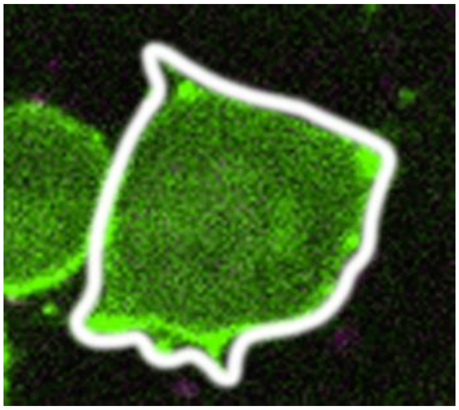

**MovieS1. Timelapse imaging of the gonialblast to 2CC TA division in control testes.** Live imaging of nos-ABD-moe::GFP, Pav::mCh tracking a gonialblast to 2CC mitosis. F-actin at the 2CC new RC is retained from the CR pool. Images taken every 15min. Each time point is a stack of 1-6 z planes. Entire time lapse is 24 hours.

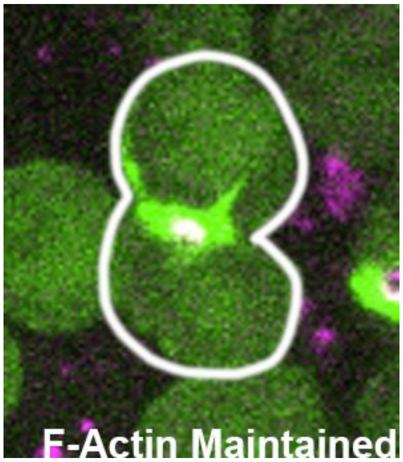


**MovieS2. Timelapse imaging of the 2CC to 4CC TA division in control testes.** Live imaging of nos-ABD-moe::GFP, Pav::mCh tracking a 2CC to 4CC mitosis. F-actin at the new RCs is retained from the CR pool, while old RCs disassemble F-actin at mitotic entry and reassemble F-actin 45 minutes after mitotic exit. Images taken every 15min. Each time point is a stack of 1-6 z planes. Entire time lapse is 24 hours.

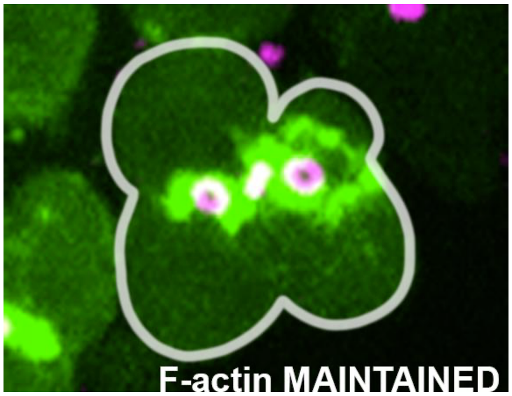


**MovieS3. Timelapse imaging of the 4CC to 8CC TA division in control testes.** Live imaging of nos-ABD-moe::GFP, Pav::mCh tracking a 4CC to 8CC mitosis. F-actin at the new RCs is retained from the CR pool, while old RCs disassemble F-actin at mitotic entry and reassemble F-actin 45 minutes after mitotic exit. Images taken every 15min. Each time point is a stack of 1-6 z planes. Entire time lapse is 24 hours.

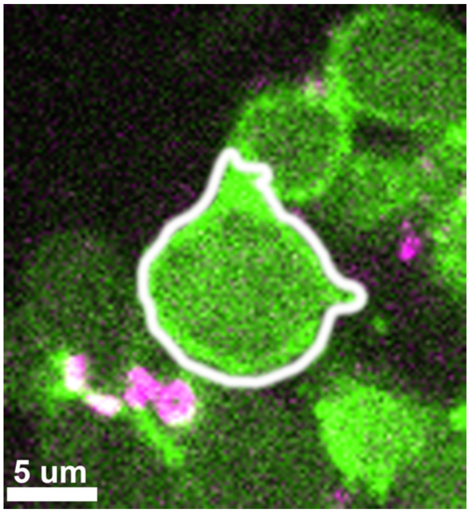


**MovieS4. Timelapse imaging of abscission in a 4CC in nos>*htl* RNAi.** Live imaging of nos-ABD-moe::GFP, Pav::mCh tracking *htl* RNAi gonia through 2CC and 4CC mitoses. At the 4CC stage, F-actin is not stably maintained and eventually incomplete cytokinesis fails, with the oldest RC executing abscission. Images taken every 15min. Each time point is a stack of 1-6 z planes. Entire time lapse is 24 hours.
